# Processing of the hepatitis E virus polyprotein can be mediated by a cellular protease

**DOI:** 10.1101/2022.12.05.519104

**Authors:** Danielle M. Pierce, Frazer J.T. Buchanan, Abigail Cox, Khadijah Abualsaoud, Joseph C. Ward, Mark Harris, Nicola J. Stonehouse, Morgan R. Herod

## Abstract

The genomes of positive-sense RNA viruses encode polyproteins that are essential for controlling viral replication. These viral polyproteins must undergo proteolysis (also termed polyprotein processing) to generate functional protein units. This proteolysis can be performed by virally-encoded proteases as well as host cellular proteases, and is generally believed to be a key step in regulating viral replication. Hepatitis E virus (HEV), a leading cause of acute viral hepatitis, translates its positive-sense RNA genome to generate a polyprotein, termed pORF1, which is necessary and sufficient for viral genome replication. However, the mechanism of polyprotein processing in HEV remains to be determined. In this study, we aimed to understand processing of this polyprotein and its role in viral replication using a combination of *in vitro* translation experiments and HEV sub-genomic replicons.

Our data suggest no evidence for a virally-encoded protease or auto-proteolytic activity as *in vitro* translation predominantly generates unprocessed viral polyprotein precursors. However, seven cleavage sites within the polyprotein (suggested by bioinformatic analysis) are susceptible to the host cellular protease, thrombin. Using a sub-genomic replicon system, we demonstrate that mutagenesis of these sites prevents replication, as does pharmacological inhibition of serine proteases. Overall, our data supports a model where HEV uses host proteases to support its replication and could have uniquely evolved not to rely on a virally-encoded protease for replication.

**Author summary:** Positive-strand RNA viruses produce polyproteins that are cleaved by proteases that control viral replication. The polyproteins of all well studied positive-strand viruses undergo proteolysis in a highly controlled manner to generate functional proteins and regulate the transition from translation to RNA replication. Proteolysis of viral polyproteins is generally performed by virally-encoded proteases, although host cell proteases are used by some viruses. In this report, we provide evidence that suggests that hepatitis E virus, a medically important human pathogen, does not encode a protease and unlike other viral polyproteins cannot undergo auto-catalytic processing. Instead, we provide evidence that the polyprotein is susceptible to proteolysis by host cell proteases and that this is essential for viral replication. Our data contradict the previous dogma of positive-sense viral replication and suggests a model where this virus has evolved to use a host protease to control viral replication and tropism.

## Introduction

Hepatitis E virus (HEV) is a leading cause of acute viral hepatitis [1, 2]. An estimated 20 million cases each year contribute to >3% of all virally related hepatitis mortalities. Human HEV is a member of the *Orthohepevirus* genus, within the *Hepeviridae* family, and is classified into 4 species groups (A-D). The genus is also independently sub-classified into eight genotypes (G1 – G8), which are found in a wide range of animals [3–5]. G1 and G2 viruses appear to be obligate human pathogens that are transmitted between humans faecal-orally, with the potential to cause large outbreaks. Viruses in G3 and G4 have been isolated in several animal species and are believed to be zoonotically transmitted to humans [6–8]. These viruses are of particular concern and have been suggested to exist within a reservoir of domestic pigs where they can be transmitted to humans, for example via poorly prepared pork products [9]. HEV infection is usually self-limiting, however, infection during pregnancy can give rise to significant mortality of up to 30% [10]. A HEV vaccine is currently only approved in China, with other treatment options including ribavirin and PEG-α-interferon [11]. This virus is therefore not only a significant global healthcare problem but also imposes risks to the farming industries and food chain security.

HEV is a positive-sense single-stranded RNA virus. The genome contains three open reading frames (ORF). ORF1 is translated into the viral polyprotein (pORF1) that is necessary and sufficient for viral RNA replication. The second and third open reading frames, ORF2 and ORF3, are translated into the viral capsid protein and a small membrane protein involved in virus release, respectively. An additional fourth open reading frame, ORF4, has also been identified in G1 viruses. Replication of the viral genome is mediated by the pORF1 polyprotein. Through sequence homology to related virus families, such as the caliciviruses and togaviruses, pORF1 has been predicted to contain at least six distinct protein domains. At the N-terminus of the polyprotein is a methyltransferase (MeT) domain, followed by the Y domain and the putative cysteine protease (PCP). In the centre of the polyprotein is a region of high sequence diversity, termed the hyper-variable region (HVR), followed by a macrodomain or X region that can bind ADP-ribose [12]. At the C-terminus of the polyprotein the domains are termed helicase (Hel) and RNA-dependent RNA-polymerase (RdRp). Based on considerable sequence homology, the functions of the MeT, X, Hel and RdRp, are highly probable. However, only the MeT, Hel and X region have been formally attributed a function. Furthermore, some regions, such as the HVR, have poor sequence homology and no function has been suggested.

The polyproteins of all well-studied positive-sense RNA viruses have been shown to undergo processing to generate the functional protein subunits (sometimes called replicase or non-structural proteins). Several studies have attempted to understand if and how HEV pORF1 undergoes processing but with varying results. Studies using heterogenous expression systems have demonstrated auto-catalytic processing of pORF1, potentially mediated by the viral PCP region, to generate smaller protein fragments, but with inconsistent results [13–19]. These studies also implicate one or more cysteine residues in the PCP as important for this proteolysis. However, these data are confused by the recent structure of the PCP region derived by X-ray crystallography, which revealed a resemblance to a fatty acid binding protein. Further data suggesting the PCP region can chelate zinc, has deubiquitinase activity, and acts with the upstream Met-Y-domain also question the function of this domain [20–22]. Other studies have suggested that pORF1 cannot be processed in heterogenous systems and is not processed in transfected cells, and therefore the intact precursor is hypothesised to be functionally active [23].

In addition to auto-catalytic pORF1 processing, one investigation suggested that the host cellular protease thrombin plays a role in processing pORF1 and implicated processing at two locations within pORF1 [24]. Thrombin is synthesised in hepatocytes as the zymogen prothrombin and secreted into the blood system [25]. In coagulation, factor Xa, factor Va and phospholipid cleave the activation peptide from prothrombin to yield thrombin in the prothrombinase complex [25]. The use of host proteases to control virus processing and replication is not unique, with many viral polyproteins cleaved in places by cellular proteases. For example, the related noroviruses use caspases to control processing of the NS1/2 non-structural protein to generate individual proteins NS1 and NS2 [26, 27] and hepatitis C virus also uses signal peptide peptidase and signal peptidase for polyprotein processing [28].

The goal of this study was to understand the processing of the HEV polyprotein and its importance for viral replication. Our data suggested that, unlike other RNA viruses, the HEV pORF1 does not have any intrinsic auto-proteolytic activity and predominantly generates a ~190 kDa precursor. However, sequence alignments identified seven conserved potential thrombin cleavage sites within pORF1, six of which we demonstrated were susceptible to thrombin proteolysis. Furthermore, we demonstrate that mutagenesis of these cleavage sites and pharmacological inhibition of thrombin are able to prevent viral replication. Thus, our data support a model where hepatocyte specific host proteases are essential for HEV replication.

## Results

### The HEV pORF1 polyprotein contains multiple potential thrombin cleavage sites

There is conflicting evidence in the literature as to if and how the HEV polyprotein is processed to functional proteins. One previous investigation by Kanade *et al*, implicated that the host protease thrombin was able to process pORF1 from a G1 HEV isolate (Sar55) [24]. Using a replicon system, they demonstrated that mutation of two thrombin cleavage junctions, after amino acid position 848 and 1220 in pORF1 prevented replication, and implicated processing of the polyprotein. Both of these cleavage sites (Figure 1) have the proline-arginine residue pairs that constitute the core thrombin cleavage site and fit within the broader thrombin recognition sequence [29]. However, upon alignment of the ~1000 genotype 1-4 HEV sequences currently available, we identified an additional five potential thrombin cleavage junctions (Figure 1) that contain a core recognition sequence and would be compatible with the extended thrombin recognition sequence. Furthermore, of these five additional sites, four are highly conserved across all available HEV sequences (Figure S1), suggesting an importance for virus replication. We therefore set out to investigate if these sites are of importance for HEV replication.

**Figure 1.**
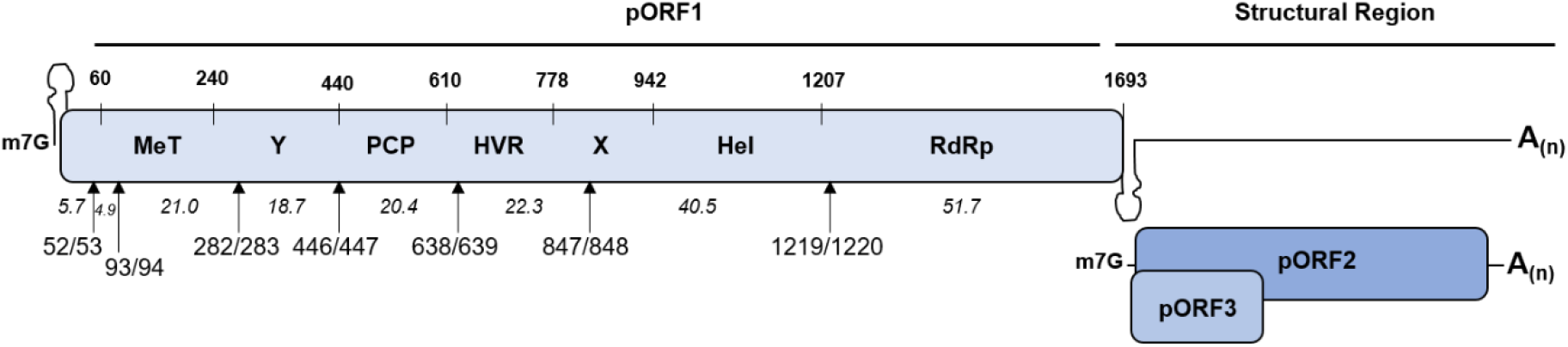
Location of the conserved pORF1 thrombin recognition sites. Schematic of the HEV genome showing the three ORF, with the pORF1 polyprotein labelled with the position of the seven predicated functional domains. The location of the conserved thrombin recognition sequences in pORF1 is indicated. The numbers in italics indicate the predicted molecular weight of products after cleavage at these positions. All numbers are the amino acid positions in the Sar55 sequence (GenBank reference AF444002).

### Thrombin proteolysis of a pORF1 C-terminal portion

Of the seven potential thrombin sites identified in our analysis, two (at PR847/848 and PR1219/1220, where the numbers refer to the amino acid position within pORF1 of the P and R residues that are essential for thrombin proteolysis) have already been shown to be susceptible to proteolysis *in vitro*, using purified enzyme and a shortened cleavage junction substrate [24]. First, it was important to establish that we could detect processing of these previously described substrates by thrombin in our assays. In contrast to the previous study, we chose to express these cleavage sites as larger fragments of pORF1. We reasoned that expression of the cleavage junctions as part of a larger polyprotein fragment would present them in a more native environment.

To this end, we generated two T7 expression constructs expressing two C-terminal portions of the genotype 1 HEV pORF1 (Sar55 isolate) polyprotein. The first portion expressed amino acids 713-1693 and contains both cleavage junctions at PR847/848 and PR1219/1220, which have been previously suggested to be thrombin substrates. The second portion contained amino acids 993-1993 and thus only the cleavage junction at PR1219/1220. Both of these constructs were used for *in vitro* coupled transcription and translation pulse chase experiments to allow the detection of both final products and any processing intermediates. To a duplicate set of reactions purified human thrombin was added 20 minutes after the start of the chase. Protein samples were taken at regular intervals, separated by SDS-PAGE, and analysed by autoradiography (Figure 2).

**Figure 2.**
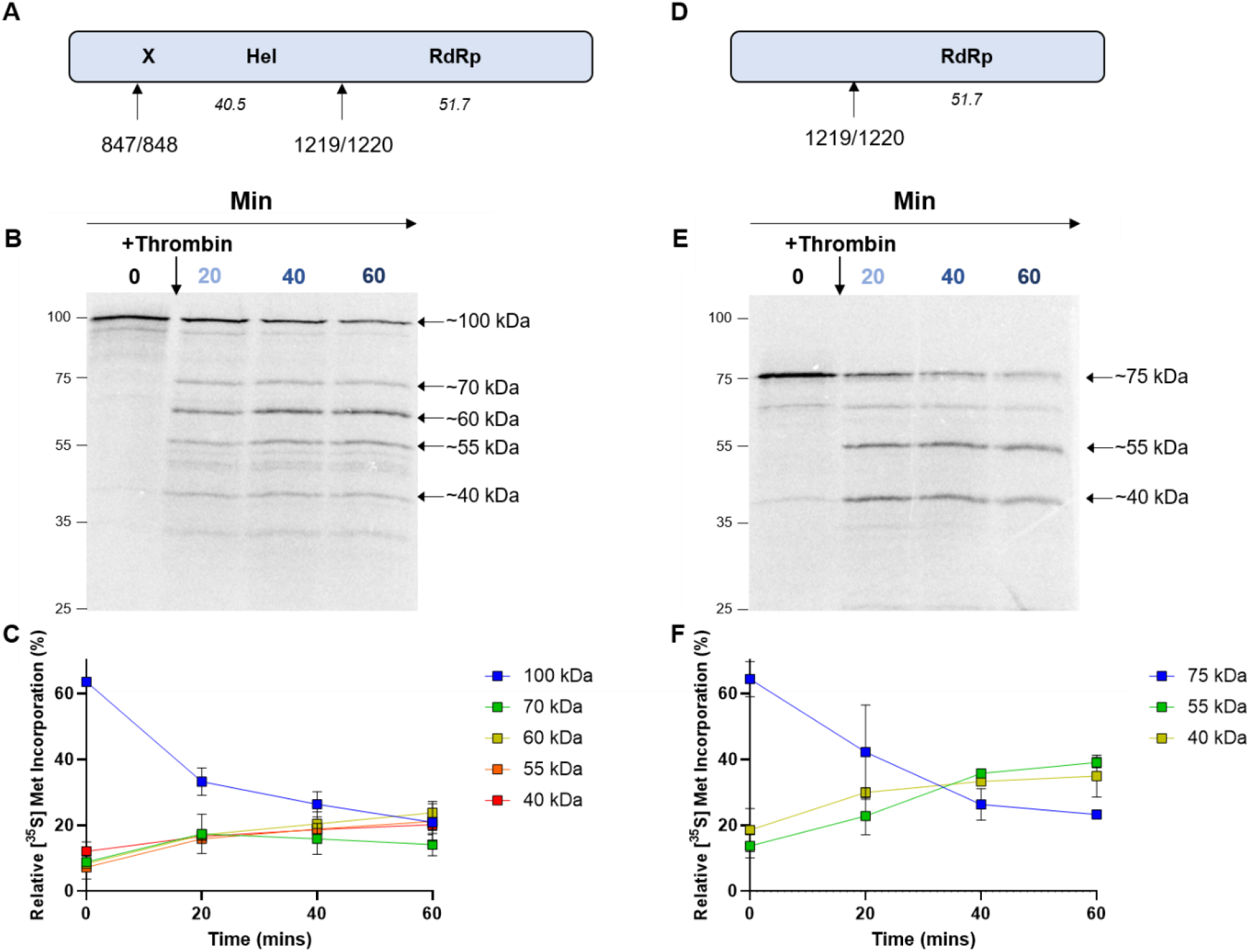
Thrombin proteolysis of the C-terminal portion of pORF1. **(A)** and **(D)** Schematics of the truncated pORF1 polyprotein expression construct in used in **(B)** and **(E).** Plasmids expressing C-terminal portions of the pORF1 polyprotein were used to template *in vitro* coupled transcription/translation reactions labelled with [^35^S] methionine before the addition of 0.5 IU of thrombin. Protein samples were taken at the indicated time-points, stopped by the addition of Laemmli buffer, proteins separated by SDS-PAGE and visualised by autoradiography. The approximate molecular weight of each product is indicated together with the molecular weight ladder in kDa on the left of each gel. The proportion of each product in **(B)** and **(E)** was quantified as a percentage of total [^35^S] incorporation and is shown in **(C)** and **(F),** respectively (n = 2 +/-SD).

In the absence of thrombin, both truncated precursors predominately generated one full-length precursor (of ~100 kDa or ~75 kDa for the 713-1693 or 993-1693 fragments, respectively), consistent with the idea that this portion of the polyprotein contains no protease activity. Upon the addition of thrombin, the 713-1693 fragment was processed into four products of ~70, ~60, ~55 and ~40 kDa. The products at ~60, ~55 and ~40 kDa increased in abundance and would correspond well to the predicted molecular weights after cleavage at either PR1219/1220 alone or at both PR847/848 and PR1219/1220. The unidentified product at ~70 kDa was less abundant and decreased in intensity over the time course. Upon the addition of thrombin, the 993-1693 portion was processed to generate a product at ~ 55 kDa. It therefore seems likely that for processing of both the 713-1693 and 993-1693 portions the ~55 kDa product is the result of processing at the PR1219/1220 predicted thrombin cleavage junction. The 993-1693 fragment also generated a product that we could not identify at ~40 kDa.

### Thrombin proteolysis of the N-terminal pORF1 portion

After analysing processing of the C-terminal portion of pORF1, we turned our attention to the N-terminal portion. This portion contains five predicted thrombin cleavage junctions, therefore processing is likely to be more complex. A T7 expression construct was generated expressing amino acids 1-712 which contained all five predicted thrombin cleavage junctions. We analysed this N-terminal region of the polyprotein using *in vitro* coupled transcription and translation in the presence or absence of thrombin (Figure 3).

**Figure 3.**
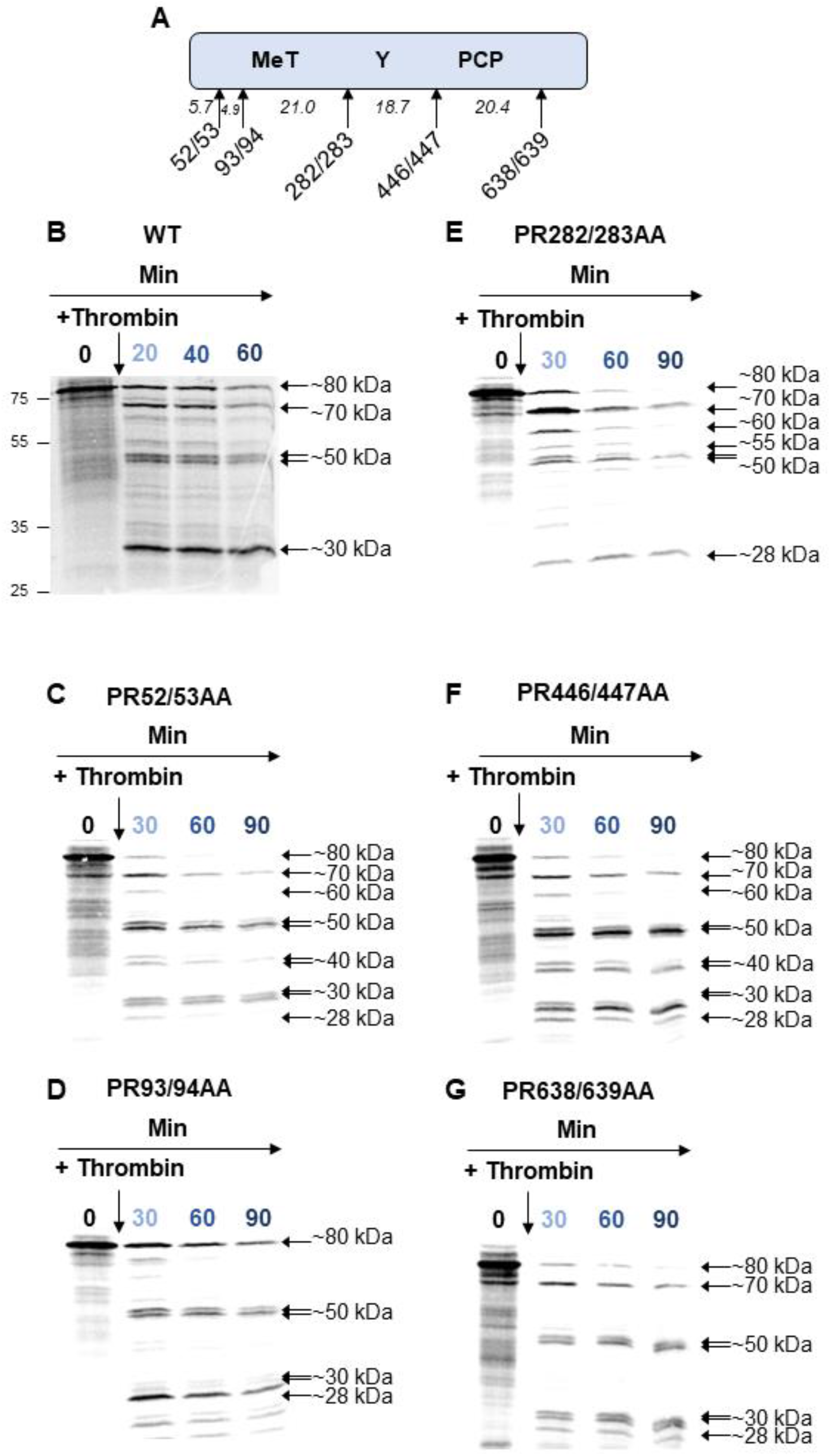
Thrombin proteolysis of the N-terminal portion of pORF1. **(A)** Schematic of the truncated pORF1 expression plasmid. **(B)** A plasmid expressing the N-terminal portion of the WT pORF1 polyprotein were used to template *in vitro* coupled transcription/translation reactions labelled with [^35^S] methionine before the addition of 0.5 IU of thrombin. **(C-G)** Plasmid expressing amino acids 1-712 of pORF1 with the indicated alanine substitutions at amino acids **(C)** PR52/53, **(D)** PR93/94, **(E)** PR282/283, **(F)** PR446/447, **(G)** PR638/639, before being used to template [^35^S] methionine labelled *in vitro* coupled transcription/translation reactions before the addition of 0.5 IU of thrombin. Protein samples were taken at indicated time-points, stopped by the addition of Laemmli buffer, proteins separated by SDS-PAGE and visualised by autoradiography. The approximate molecular weight of each product is indicated together with the molecular weight ladder on the left of the gel.

Firstly, before the addition of thrombin this polyprotein was visualised as a single protein that is approximately the predicted molecular weight of the uncleaved product. Therefore, in common with the data above for the C-terminal region, the N-terminal portion of the polyprotein was also unable to undergo significant auto-catalytic proteolysis, despite including the putative viral protease (PCP).

Upon the addition of thrombin, the protein underwent proteolysis to produce at least five new products. Of these new products the largest, of ~70 kDa, seems likely to result from cleavage of ~10 kDa from the N-terminus of the polyprotein, which is consistent with processing at the predicted PR93/94 site. Cleavage of the construct generated products at ~30 kDa and ~55 kDa, suggesting these products have N- and C-termini within the first 712 amino acids of pORF1. The generation of these products suggest at least some, if not all, of the predicted sites at amino acids PR52/53, PR282/283, PR446/447 and PR638/639 are susceptible to thrombin proteolysis. Using the predicted molecular weight of the different cleavage products, it seems likely that the ~30 kDa product is the result of processing at PR282/283, and the ~ 55kDa product is the result of processing at PR446/447. However, these products could have arisen through multiple combinations of proteolysis events. For example, the ~30 kDa product could be the result of proteolysis at both the PR93/94 and PR446/447 sites simultaneously. It was therefore still difficult to establish whether all of the predicted sites were cleaved by thrombin from the data generated in these assays alone.

### Mutagenesis of thrombin proteolysis sites helps elucidate cleavage pathways

To help elucidate these possible cleavages further we used site directed mutagenesis to introduce alanine substitutions at either the PR52/53, PR93/94, PR282/283, PR446/447 or PR638/639 residue pairs within the context of the 1-712 precursor. These substitutions would be expected to prevent recognition and proteolysis by thrombin. This generated a new series of T7 expression constructs in which each protease site was removed. This new series of constructs were used for *in vitro* transcription and translation in the presence of thrombin as above (Figure 3). To understand the effect of each mutation of processing, the relative proportion of each product was quantified and compared to the wild-type (WT) control. For ease of interpretation, the main differences between the different constructs were plotted in comparison to WT (Figure 4). Importantly, if we removed a genuine thrombin cleavage site, we would anticipate that the products formed by proteolysis would be different to the WT control (i.e. a disappearance of smaller products and concomitant accumulation of larger protein products). However, if the site was not genuinely susceptible to thrombin mediated proteolysis, then we would expect to see no difference in the pattern of proteolysis compared to the WT control.

**Figure 4.**
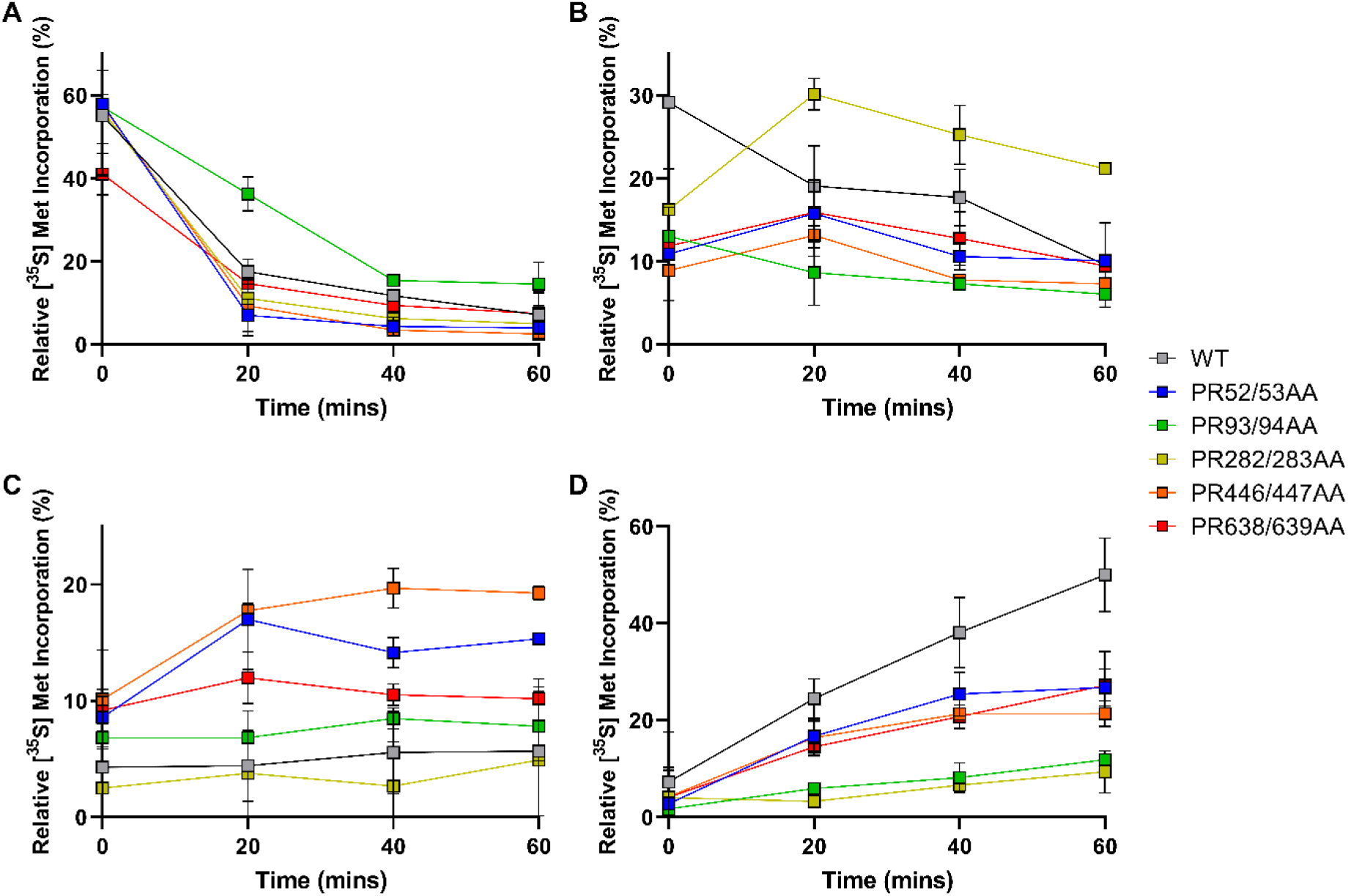
Thrombin proteolysis of the N-terminal portion of pORF1. Site directed mutagenesis was used to introduce alanine substitutions at either the PR53/54, PR93/94, PR282/283, PR446/447 or PR638/639 residue pairs within the context of the 1-712 precursor. These plasmids were used to template *in vitro* coupled transcription/translation reactions labelled with [^35^S] methionine before the addition of 0.5 IU of thrombin. Proteins were separated by SDS-PAGE and visualised by autoradiography (shown in Figure 3). The relative proportions of the **(A)** ~80 kDa, **(B)** ~70 kDa, **(C)** ~40 kDa and **(D)** ~30 kDa proteins were quantified from each of these substitutions in comparison to the WT control (n = 2 +/-SD).

Mutation of the PR93/94 position decreased the rate of processing of the ~80 kDa (i.e. full-length) protein, and slowed the appearance of products at ~30 kDa and ~70 kDa. These observations therefore suggest that processing at the PR93/94 position can take place and is key for generating the ~70 kDa product.

Mutation at the PR282/283 position slowed the appearance of the ~30 kDa product, as well as generating a novel product at ~55 kDa, not observed with any of the other constructs. These data suggest processing at the PR282/283 site can occur and is important for generating the ~30 kDa product.

The PR446/447AA mutation increased the appearance of the ~40 kDa product, and moderately changed the appearance of the ~30 kDa product. Thus, suggesting processing can occur at this position, and the ~40 kDa product is the result of cleavage at PR282/283 and possibly at PR638/639.

Similar changes were observed with either the PR52/53AA or PR638/639AA mutations in comparison to the WT but these were less pronounced. This could either be due to a lack of efficient cleavage at these positions, or the resultant differences being too small to visualise in these assays. Taken together, these data provide evidence for processing at five predicted thrombin cleavage sites, PR93/94, PR282/283, PR446/447, PR847/847 and PR1219/1220.

### *In vitro* proteolysis of pORF1

Our experiments could not detect auto-catalytic activity of pORF1 fragments in an *in vitro* transcription and translation system, in contrast to many other viruses [30, 31]. To verify that no auto-catalytic processing was also observed when full-length pORF1 was expressed, a T7 expression construct was generated expressing the entire pORF1 coding sequence (using the same genotype 1 Sar55 isolate). As a control to confirm that auto-catalytic processing is possible in this system we generated an equivalent construct expressing the non-structural ORF1 polyprotein from murine norovirus (MNV). This viral polyprotein was chosen based on its known processing profile, similar layout of functional domains and similar polyprotein length [26, 32, 33]. These two expression constructs were used to template *in vitro* coupled transcription and translation, labelled with [^35^S] methionine. As before, samples were taken at regular time points, proteins separated by SDS-PAGE and analysed by autoradiography (Figure 5).

**Figure 5.**
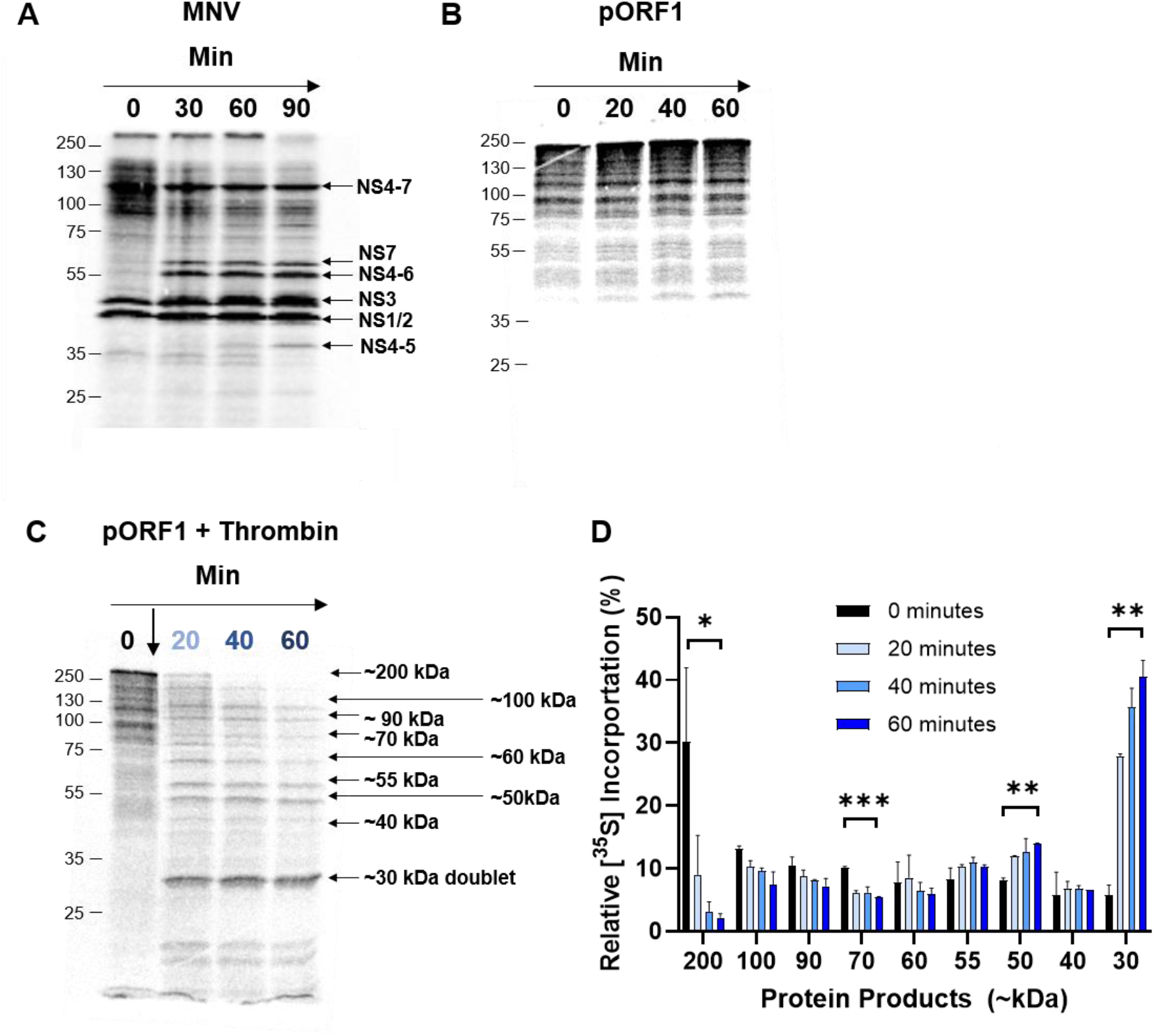
Thrombin proteolysis of pORF1. Plasmids expressing MNV polyprotein **(A)** or the HEV pORF1 **(B)** were used to template *in vitro* coupled transcription/translation reactions labelled with [^35^S] methionine. Samples were taken at regular intervals, reactions stopped by the addition of Laemmli buffer, proteins separated by SDS-PAGE and visualised by autoradiography. The identity of MNV products is indicated together with the molecular weight ladder on the left of each gel. **(C)** To a duplicate HEV pORF1 reaction, 0.5 IU thrombin was added as indicated and product of thrombin proteolysis indicated together with their approximate molecular weight. **(D)** The relative portion of each product was quantified as a percentage of the total [^35^S] incorporation (n = 2 +/-SD; *=p<0.05, **=p<0.01, ***=p<0.001).

With the MNV control plasmid, at least six distinct protein products could be observed which correlated well to a range of different mature and precursor protein products that would occur from auto-catalytic proteolysis (Figure 5A) [26, 32–34]. These data demonstrate that auto-catalytic processing is possible in this *in vitro* system.

In contrast, translation of pORF1 produced predominantly just one product of approximately the predicted size for full-length unprocessed HEV pORF1 (Figure 5B). There was a smaller abundance of lower molecular weight products produced that we attribute to early termination events, which is common when performing *in vitro* translation of large proteins [35]. These results suggested that full-length pORF1 has no intrinsic auto-catalytic activity. To stimulate any intrinsic protease activity of pORF1, we titrated in divalent metal ions or fatty acids, both of which have been suggested to interact with the PCP domain [22]. We also attempted to change the oxidation state of the reactions with reducing agents and added viral RNA in an attempt to induce auto-catalytic proteolysis. However, none of these approaches stimulated proteolysis in this assay (data not shown). Finally, we investigated the products formed from thrombin mediated proteolysis of full-length pORF1 (Figure 5C). Upon the addition of exogenous thrombin, pORF1 underwent proteolysis to generate at least nine distinct additional protein products, each of which were quantified (Figure 5D). There was also a clear decrease in the full-length pORF1 upon the addition of thrombin. Larger molecular weight products, at ~100, ~90 and ~70 kDa decreased in relative abundance over the duration of the experiment. The relative abundance of products at ~60 kDa and ~40 kDa did not significantly change. There was a small increase in the abundance of the product at ~55kDa, although it was not significant over the duration of the experiment, and a clear increase in the abundance of the smaller protein products at ~50 kDa and the doublet at ~30 kDa.

### Immunoprecipitation of polyprotein products

A notable observation from our *in vitro* processing data was that there were more than the seven cleavage products that would be predicted from full proteolysis. We hypothesised therefore that some of these additional products were protein intermediates. Therefore, we sought to identify the composition of some of these potential precursors in addition to fully processed proteins. Due to the lack of suitable antibody reagents for immunoprecipitation, we adapted the pORF1 T7 expression vector by incorporating a HA-tag at the C-terminus of pORF1 which would allow immunoprecipitation of RdRp containing products and precursors. This expression vector was used in an *in vitro* coupled transcription and translation assay with [^35^S] methionine before HA-containing products were immunoprecipitated, separated by SDS-PAGE and imaged by autoradiography (Figure 6).

**Figure 6.**
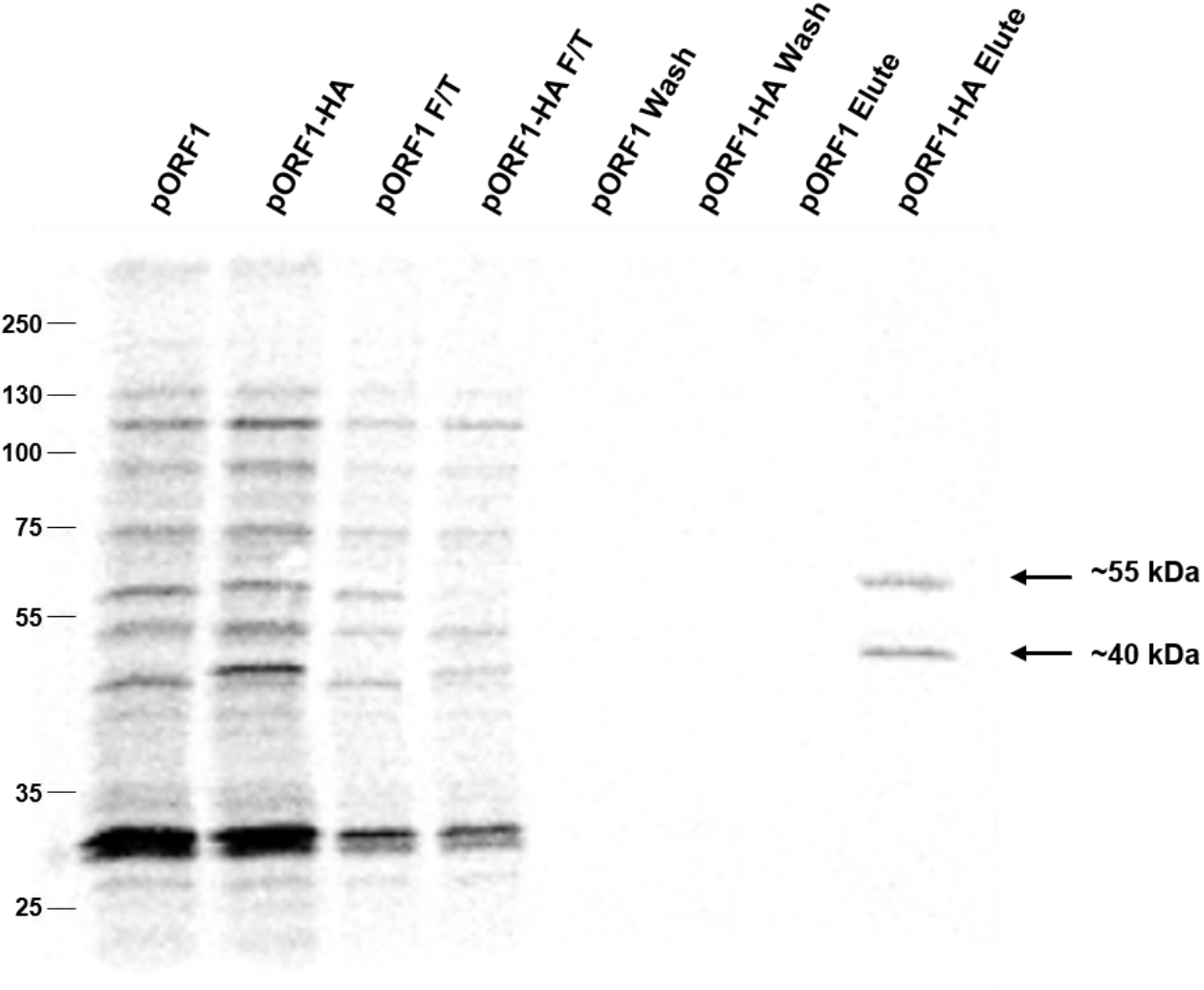
Immunoprecipitation of HEV pORF1 products. Plasmids expressing HEV pORF1 or pORF1-HA containing a C-terminal HA-tag were used to template [^35^S] Met labelled pulse-chase *in vitro* coupled transcription/translation reactions. Reactions were incubated with thrombin for 90 minutes before proteins were immunoprecipitated with anti-HA antibody. The pre-IP samples, flow through (F/T), wash and elute samples were separated by SDS-PAGE and visualised by autoradiography. The immunoprecipitated ~40 kDa and ~55 kDa products are indicated together with the molecular weight ladder on the left of the gel. Representative result from one of three experiments.

Thrombin-mediated processing of HA-labelled pORF1 yielded at least nine additional products, which are approximately equivalent to the untagged pORF1 plasmid and similar to the products observed in earlier pulse-chase experiments. Notably, some products in the pORF1-HA sample appeared to have a marginally greater molecular weight compared to the unlabelled pORF1 sample, as may be anticipated with a C-terminal epitope extension. Immunoprecipitation of the HA-tagged sample yielded two clear products of ~40 kDa and ~55 kDa. It seems likely the ~55 kDa product corresponds to the C-terminal fragment generated from cleavage at the PR1219/1220 position, thus corresponding to the RdRp domain. The smaller ~40 kDa product would be consistent with the results in Figure 2 and could be the result of inadvertent off-target proteolysis, incorrect translation initiation or a co-immunoprecipitated product.

### Preventing thrombin proteolysis inhibits viral replication

To understand if any of these sites are important for virus replication we utilised a sub-genomic replicon (SGR) of the G1 Sar55 HEV sequence, which contained a nano-luciferase (nLuc) reporter sequence in place of the viral structural proteins. Monitoring the production of luciferase allows for indirect quantification of virus replication. This SGR was modified by alanine substitution of each proline arginine pair as above, to prevent recognition and proteolysis by thrombin. This generated seven new replicons each with an individual proline arginine amino acid pair mutated (termed PR52/53AA, etc). RNA from these replicons were transfected into Huh7 cells along with a wild-type (WT) control replicon or a replicon containing an inactivating mutation in the RdRp active site (GNN), and luciferase activity was monitored over four days to measure RNA replication (Figure 7A and Figure S2).

**Figure 7.**
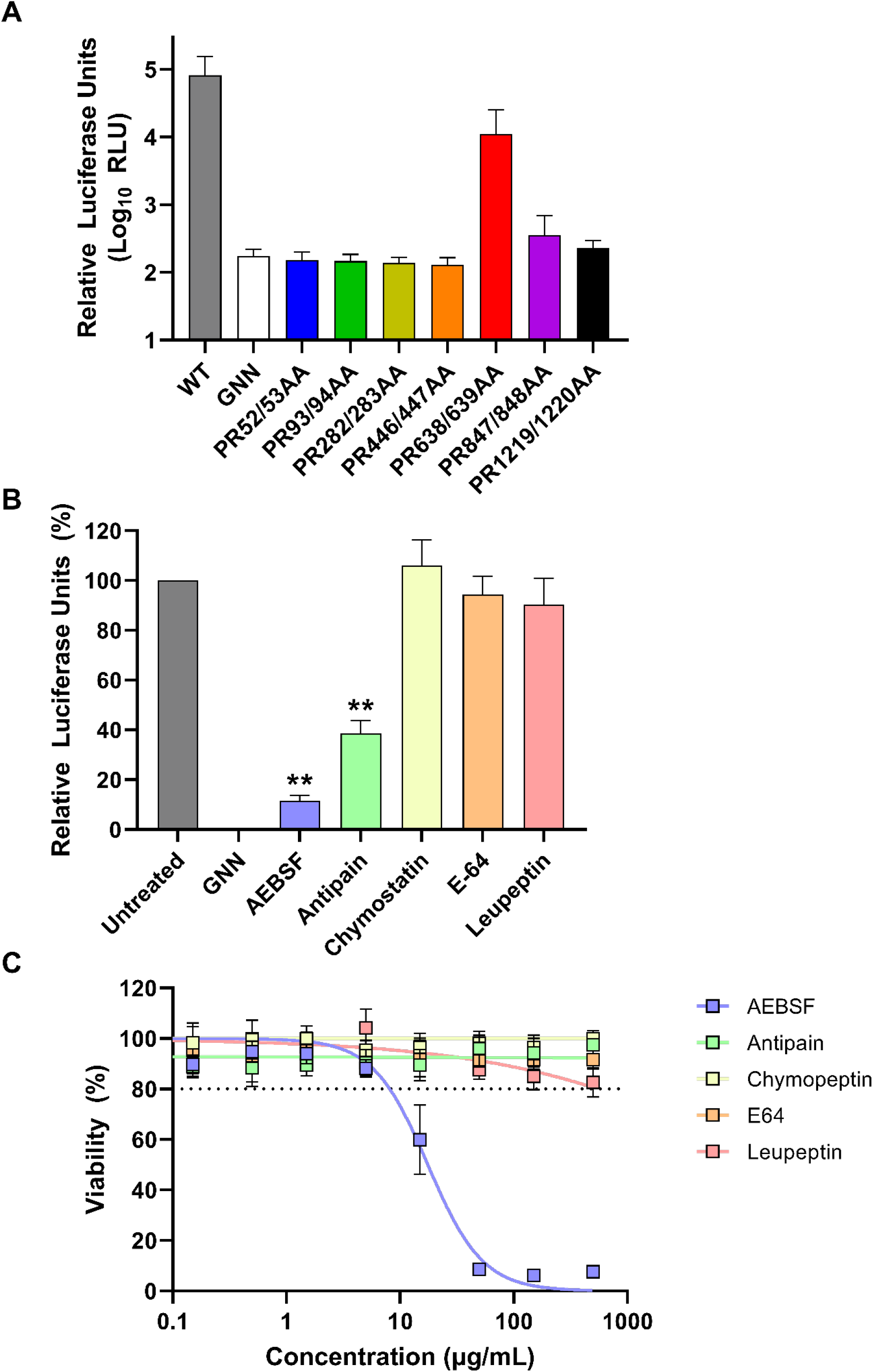
Preventing thrombin proteolysis prevents HEV replication. **(A)** Huh7 cells were electroporated with HEV replicon RNA containing the indicated mutations at predicted thrombin cleavage junctions, in addition to the WT and GNN control replicons. Cells were harvested at 96 h post-electroporation and luciferase activity determined. Data shown represents log_10_ of mean relative luciferase activity at 96 h post-electroporation (n = 3 +/-SEM). **(B)** Huh7 cells were electroporated with the WT HEV replicon RNA or GNN control replicon before the addition of AEBSF (2.5 μg / mL), antipain (50 μg / mL), chymostatin (25 μg / mL), E-64 (100 μg / mL) or leupeptin (175 μg / mL) at 24 h post-electroporation. Cells were harvested at 96 h post-electroporation and luciferase activity determined. Data shown represents mean relative luciferase activity at 96 h post-electroporation normalised to the untreated control (n = 3 +/-SEM, *=p<0.05, **=p<0.01 compared to WT). **(C)** Huh7 cells were incubated with a serial dilution of protease inhibitors for 72 h before cell viability was measured by MTS assay. Data are expressed as mean percentage cell viability normalized to untreated controls (n = 3 +/-SEM).The WT HEV replicon gave a >100-fold increase in luciferase activity over the duration of the experiment compared to the replication defective (GNN) control. In contrast, transfection of all but one of the seven replicons with proline-arginine substitution significantly impaired replication, with luciferase activity equivalent to the replication-defective control (GNN). The exception was PR638/639AA which demonstrated an approximate 2-fold reduction in luciferase activity compared to the WT replicon, although this reduction was not statistically significant. These data would suggest that all but one of the PR residues where thrombin is predicted to cleave are essential for viral replication.

### Inhibition of serine proteases prevents HEV replication

The HEV PCP has been suggested to have a cysteine active site or act like a metalloprotease [17–19, 22, 36]. In contrast, thrombin is a serine protease. These differences allowed us to use a range of commercially available selective protease inhibitors to test inhibition of replicon replication. Huh7 cells were therefore transfected with the WT replicon or GNN control prior to the addition of a range of protease inhibitors at 24 h post-transfection and monitoring of luciferase activity over four days to measure RNA replication (Figure 7B and Figure S2).

Five protease inhibitors were chosen based on their specificity. AEBSF is an irreversible serine protease inhibitor that inhibits chymotrypsin-like proteases including trypsin and thrombin. E-64 is an irreversible cystine protease inhibitor that includes papain-like proteases. Leupeptin is an inhibitor which can target a range of proteases including cysteine, serine and threonine proteases, including trypsin and papain, but importantly has lower specificity for thrombin. Antipain is a reversible serine/cysteine protease inhibitor of broad spectrum with a similar action to leupeptin, but which includes thrombin. Chymostatin is an inhibitor of many proteases, including chymotrypsin and papain as well as chymotrypsin-like serine proteinases. A single concentration of each inhibitor was chosen based on the maximal tolerated concentration.

Upon treatment with this range of inhibitors, only AEBSF and antipain significantly inhibited replicon replication, reducing luciferase activity ~80% and ~60% at 4 days post-electroporation, respectively. The other inhibitors tested did not change luciferase activity compared to the untreated control.

To determine if any of the reduced replication was the result of cytotoxicity, Huh7 cells were incubated with a serial dilution of the protease inhibitors used and cell viability measured by MTS assay (Figure 7C). AEBSF was the only protease inhibitor to display cytotoxicity at any of the concentrations tested, with a CC_50_ of ~13 μg/mL. However, no cytotoxicity was observed at 2.5 μg/mL, which was used in the HEV replication assays. Taken together, the results from the replicon replication assays suggest thrombin or another serine protease is an important host factor for HEV replication.

## Discussion

Positive-sense RNA viruses in general encode polyproteins that undergo precise proteolysis to generate functional units referred to as non-structural (NS) proteins. These NS proteins assemble into active genome replication complexes also termed the replicase. Insight into viral polyprotein processing is therefore important for understanding how the replication complex is formed and the functional proteins within this.

Multiple studies have attempted to understand processing of the HEV polyprotein (pORF1), providing some data or suggestions on how this may occur. The results from these studies can be divided into two broadly conflicting models. Several reports have provided evidence for pORF1 processing, generating either two larger products of ~110 and ~80 kDa or multiple products ranging in size from ~18 to ~120 kDa [14–19]. In contrast, data from several studies suggest that pORF1 is not processed and could function as a single polyprotein [23, 37, 38]. This is supported by the observations that no protease activity has been attributed to the postulated viral protease, PCP [20, 22]. The reasons behind the contradictions in these studies are not clear but is possibly due to the wide range of expression systems (i.e. *in vitro* transcription/translation, insect cells, vaccinia expression, and various mammalian cell lines) and methods (i.e. western blot with custom generated antibodies and/or radiolabelling) used for detection. Here, we found that the pORF1 polyprotein was unable to be processed auto-catalytically using a well-established *in vitro* transcription and translation system. We, and others, have used this system to study the processing of many positive-sense RNA viruses. For example, MNV ORF1 can undergo auto-catalytic processing in this system to generate the same range of proteins found during natural infection [26, 32, 33]. To attempt to stimulate intrinsic protease activity of pORF1 in this system we supplemented the reactions with various metal ions, reducing agents, fatty acids and cellular extracts. Despite this, none of these factors could elicit protease activity and change the products formed. In addition, we mutated eight of the amino acids suggested to be the protease active site [17–19, 22], however again none of the mutants changes the products formed (data not shown). All these observations support the previous studies suggesting that pORF1 has no auto-catalytic activity, and that the PCP domain does not have protease activity.

In addition to this work, a single study by Kanade *et al*, implicated two host proteases, thrombin and factor Xa, as important for HEV replication [24]. They showed that these enzymes were able to cleave the purified fragments of pORF1 at two places and siRNA silencing of thrombin expression reduced HEV replication. The authors therefore suggested that thrombin was important for HEV replication. By comparative alignment of all currently available HEV sequences we identified additional sites that match the thrombin recognition sequence [29], six of which were highly conserved (Figure S1). Indeed, we found that addition of exogenous thrombin to the *in vitro* transcription/translation assays processed the pORF1 polyprotein into at least 9 defined products. Using a combination of mutagenesis and polyprotein truncations we demonstrated that at least six of these seven sites were cleavable by thrombin *in vitro* and prevented replicon replication. Interestingly, these were sites of high sequence conservation across HEV isolates. The remaining site (PR638/639AA) was poorly conserved, only reduced replication ~2-fold when mutated, and has low homology to the thrombin cleavage consensus. These data would suggest therefore that the PR638/639AA sequence is not a genuine processing site, or at least these residues are not essential for viral genome replication. Given that we observed at least nine products (excluding full-length pORF1), yet there are only 6 cleavage sites, some of the products observed must be polyprotein precursors. To help us theoretically assign the nine observed products to different cleavage events, we compared the products from proteolysis of the full length pORF1 to the truncated regions (Figure 8). Based on molecular weight, it is likely that products at ~100 to ~60 kDa were large precursors spanning multiple domains. For example, a ~100 kDa product is possible the result of cleavage at position PR282/283 and PR1219/1220, whereas products at ~80 kDa could be result of processing at PR446/447 and PR1219/1220. Processing at PR1219/1220 would yield a ~55 kDa product that would include the viral RdRp. This product was also immunoprecipitated from the reaction using C-terminally epitope tagged constructs, supporting this identification. It is also clear that the ~50 kDa products and doublet at ~30 kDa are only present at the N-terminal fragment of the polyprotein and likely be the result of cleavage at the PR446/44 and PR282/283 positions, respectively. The larger products could undergo further proteolysis to generate many of the same final products, for example, the ~40 kDa product found within the C-terminus is likely to be the result of thrombin proteolysis at PR847/848 and PR1219/1220 and could be derived from either ~100 or ~80 kDa precursors. Further work is therefore needed to conclusively confirm the identity of all the observed products.

**Figure 8.**
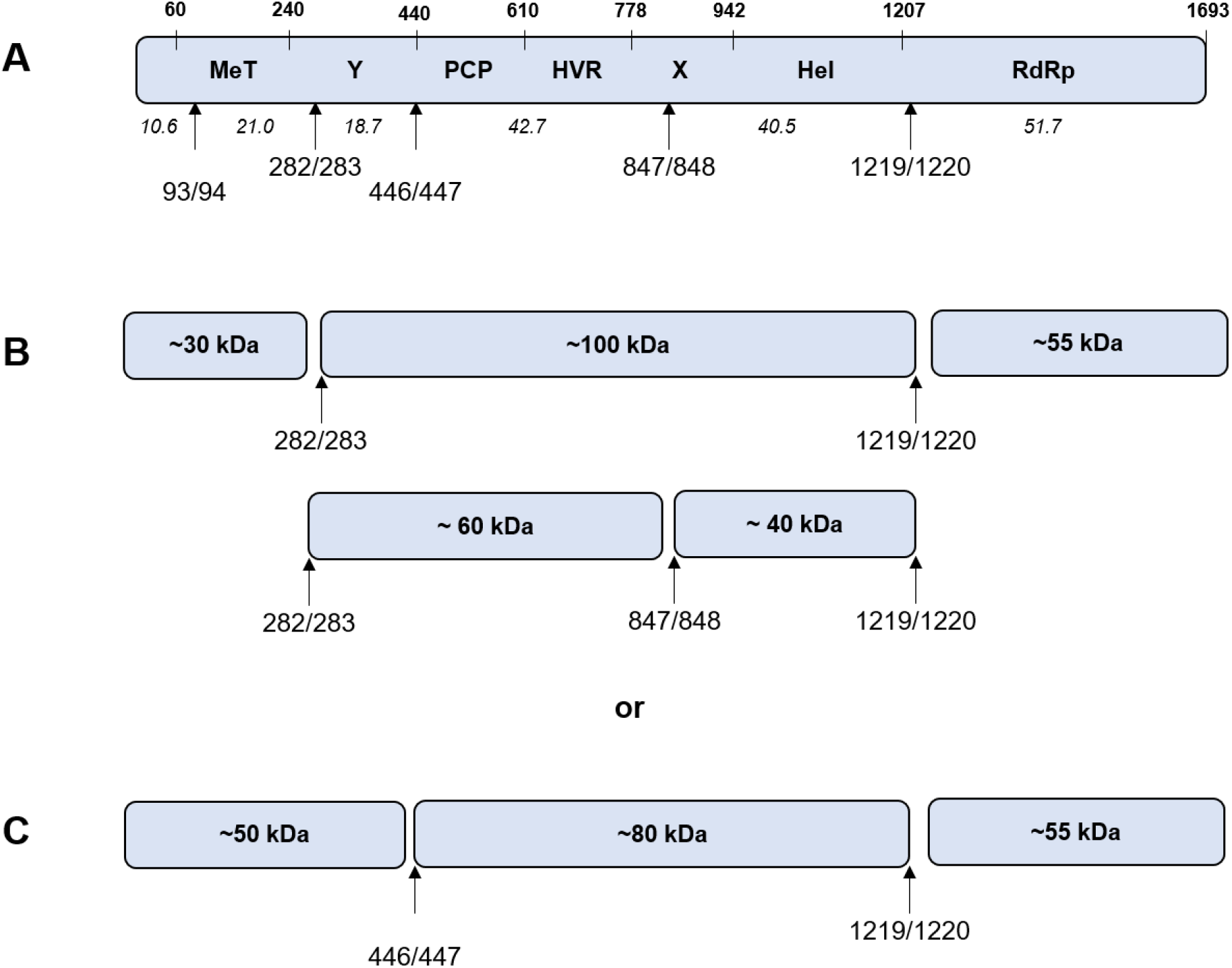
Overview of thrombin-mediated proteolysis of pORF1. **(A)** Schematic of the pORF1 showing the cleavage sites which we present data for processing by thrombin. Numbers in italics show the predicted molecular weight of products after theoretical complete thrombin proteolysis. **(B)** The observed ~ 30, ~100 and ~55 kDa products would be explained by processing at the PR282/283 and PR1219/1220 position. Subsequent processing at PR847/848 would yield ~60 and ~40 kDa products. **(C)** The observed ~ 50, ~80 and ~55 kDa products would be explained by processing at the PR446/447 and PR1219/1220 position.

Many viruses use host cellular proteases to regulate polyprotein processing or other aspects of the viral replication cycle. For example, hepatotropic viruses such as hepatitis C virus use host signal peptidases to cleave its polyprotein at crucial points [28]. It is not clear from the data presented here whether the host protease thrombin is solely responsible for HEV pORF1 processing, or it acts in conjunction with a host or viral protease. It is also possible a mechanism exists where a host protease is responsible for primary polyprotein processing which releases a viral protease to enact secondary processing. We could not find any evidence of this in our experiments. However, the development of new reagents is needed in order to investigate HEV pORF1 processing in cells before these questions can be answered more completely, or indeed the role of thrombin in pORF1 processing can be confirmed conclusively.

If thrombin is solely responsible for the processing of pORF1 it would represent a new mechanism of polyprotein control. Thrombin is synthesised specifically in hepatocytes as the inactive complex multi-domain zymogen prothrombin [39, 40]. Prothrombin consists of an N-terminal Gla domain, which is modified for membrane association in a co-translational vitamin K dependent reaction. This directly precedes two kringle domains, K1 and K2, and the main C-terminal protease domain (consisting of A and B chains). Prothrombin is secreted into the blood where the Gla and kringle domains are removed by enzymes in the prothrombinase complex to generate the active enzyme thrombin (via intermediates) in the clotting cascade [25, 41]. However, several reports have identified active thrombin at detectable levels in hepatocytes where it is believed to play some role in cancer regulation [42–44]. The enzyme could therefore be available in cells at sufficient concentrations to allow for pORF1 cleavage. Furthermore, the highly regulated tissue expression could in part account for the viral tropism. Alternatively, a serine protease that is active in an intracellular compartment with specificity similar to thrombin, e.g. Hepsin in the endoplasmic reticulum, may be responsible for pORF1 proteolysis, which would explain findings here.

If thrombin is essential for HEV processing or genome replication, this has important implications for viral zoonosis. Viruses that are able to replicate across species must have overcome host cell restriction factors. Thrombin is common to all mammals and is genetically, structurally, and functionally similar across human, bovine and porcine species [45, 46]. It could therefore be an advantage for viral transmission to rely on a key and conserved host enzyme. However, this could open up new avenues for novel therapeutic design. Work is ongoing to fully dissect the role of thrombin in HEV replication in cells, and how this can be exploited for novel therapeutic design.

## Materials and Methods

### Cell lines and plasmids

Huh7 cells were maintained in Dulbecco’s modified Eagle’s medium with glutamine (Sigma-Aldrich) supplemented with 10 % (v/v) FCS, 1 % (v/v) non-essential amino acids 50 U *I* mL penicillin and 50 μg / mL streptomycin.

Plasmid carrying wild-type HEV replicon expressing GFP, pSK-E2-GFP, was a kind gift from Dr Patrizia Farci and has been described previously [47]. This plasmid was modified to replace the GFP open reading frame with nano-luciferase as previously described [48]. Mutations within these plasmids were performed by standard two-step overlapping PCR mutagenesis. Negative control replicons were generated containing a double point mutation in the RdRp active site GDD motif (GNN).

For coupled *in vitro* transcription and translation experiments, pcDNA3.1(+) based expression plasmids were generated by PCR. Firstly, to facilitate cloning a *NotI* restriction enzyme site within the HEV pORF1 coding region was removed by silent mutagenesis. Subsequently, the relevant HEV sequence was amplified to including flanking *Not*I restriction enzymes and upstream Kozak modified translational start site. To insert a HA-epitope at the C-terminus of pORF1 the reverse PCR primer contained a HA sequence before the stop codon. The *Not*I-digested PCR products were cloned into *Not*I digested pcDNA3.1(+) (Thermo Fisher Scientific). The sequence of all primers and plasmids are available on request.

### Coupled transcription and translation reactions

Coupled *in vitro* transcription and translation assays were performed using the TNT Quick Coupled Transcription/Translation system (Promega) following manufacturer’s instruction. Reactions contained 10 μL rabbit reticulocyte lysate with 500 ng of pcDNA T7 expression plasmid and 0.5 μL [^35^S] methionine (PerkinElmer). Reactions were incubated at 30°C for 40 minutes before being chased with 2 μl of 50 mg / mL unlabelled methionine. Plasma-purified human thrombin (Merck) was then added to reactions as required from a 1 IU / μL stock. Reactions were stopped at regular intervals by the addition of 2 x Laemmli buffer. Samples were separated by SDS-PAGE before visualisation of radiolabelled products by autoradiography.

### *In vitro* transcription

The HEV replicon plasmids were linearised with *BglII* before being used to generate T7 *in vitro* transcribed RNA using the HiScribe T7 ARCA mRNA kit with tailing following manufacturer’s instructions (Promega). RNA was purified using an RNA clean and concentrate kit (Zymo Research) and the quality was checked using MOPS/formaldehyde agarose gel electrophoresis.

### Replication assays

Replicon experiments were conducted as previously described [48]. Briefly, Huh7 cells were detached by trypsin, washed twice in ice-cold DEPC-treated PBS and re-suspended at 1 x 10^7^ cells / mL in DEPC-treated PBS. Subsequently 400 μL of cells was mixed with 2 μg of RNA transcript, transferred to a 4 mm gap electroporation cuvette (SLS) and pulsed at 260 V, 25 ms pulse length in a Bio-Rad Gene Pulser (Bio-Rad). Electroporated cells were recovered into 4 mL media, seeded into replicate 6-well tissue culture vessels, and replication measured at 24 h intervals using Nano-Glo luciferase assay system (Promega). For inhibitor treatment the electroporated cells were seeded into replicate 24-well plates, allowed to adhere for 24 h before the media was replaced with fresh media containing antipain, AEBSF, leupeptin or pepstatin (all Sigma-Aldrich), at the indicated concentration. The CC_50_ experiments were conducted by seeding cells into 96-well plates, allowing to adhere for 24 h before addition of a serial dilution of protease inhibitors and measurement of cell viability 72 h later using the CellTiter AQueous One solution (Promega), following manufacturer’s instructions.

### Immunoprecipitation

Immunoprecipitation reactions were performed using Dynabeads Protein G (Invitrogen). To bind the antibody to magnetic beads, 10 μL of the anti-HA rabbit antibody (Sigma-Aldrich) was mixed with 195 μL PBS and incubated at room temperature with 50 μL magnetic beads, shaking for 1 h, after which the supernatant was removed from the beads. Transcription and translation reaction samples were mixed with 200 μL PBS and incubated shaking at room temperature with 25 μL of Dynabeads as a pre-clear step. The tube was placed on the magnet and the supernatant removed and added to the 50 μL of Dynabeads with the antibody bound and incubated at room temperature shaking for 1 h. The flow through was removed and added to 2x Laemmli buffer. The beads were washed three times with PBS with 0.02 % Tween-20 and each wash supernatant retained. Proteins were eluted from the beads by adding 50 μL of 2x Laemmli buffer and heating to 100°C.

## Author contributions

MRH, NJS and MH designed the study and wrote the manuscript. DMP and AC conduced the *in vitro* translation experiments. FJTB and JCW conducted the replication assays. KA conducted the survival assays. DMP, FJTB, JCW and MRH analysed the data. MRH, NJS and MH provided supervision.

## Acknowledgements

We thank Patrizia Farci (National Institute of Allergy and Infectious Diseases, Bethesda) for the genotype 1 HEV replicon.

## Materials & correspondence

Correspondence and materials requests should be directed to MRH.

