## Supplementary material for "Processing of the hepatitis E virus polyprotein can be mediated by a cellular protease": Figure S1

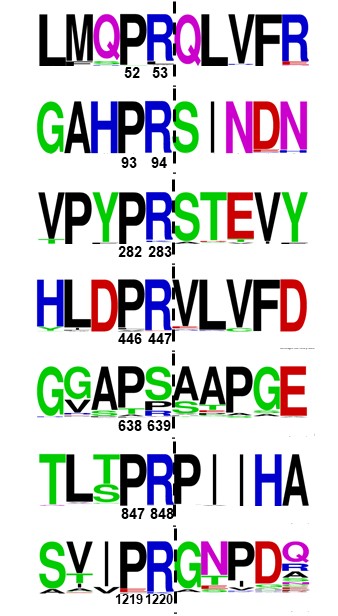

**Figure S1. Alignment of the conserved pORF1 thrombin recognition sites.** Sequence logos show the conservation of amino acids at the predicted thrombin cleavage junctions. All numbers are the amino acid positions in the Sar55 sequence (GenBank references AF444002).
