## Supplementary material for "Processing of the hepatitis E virus polyprotein can be mediated by a cellular protease": Figure S2

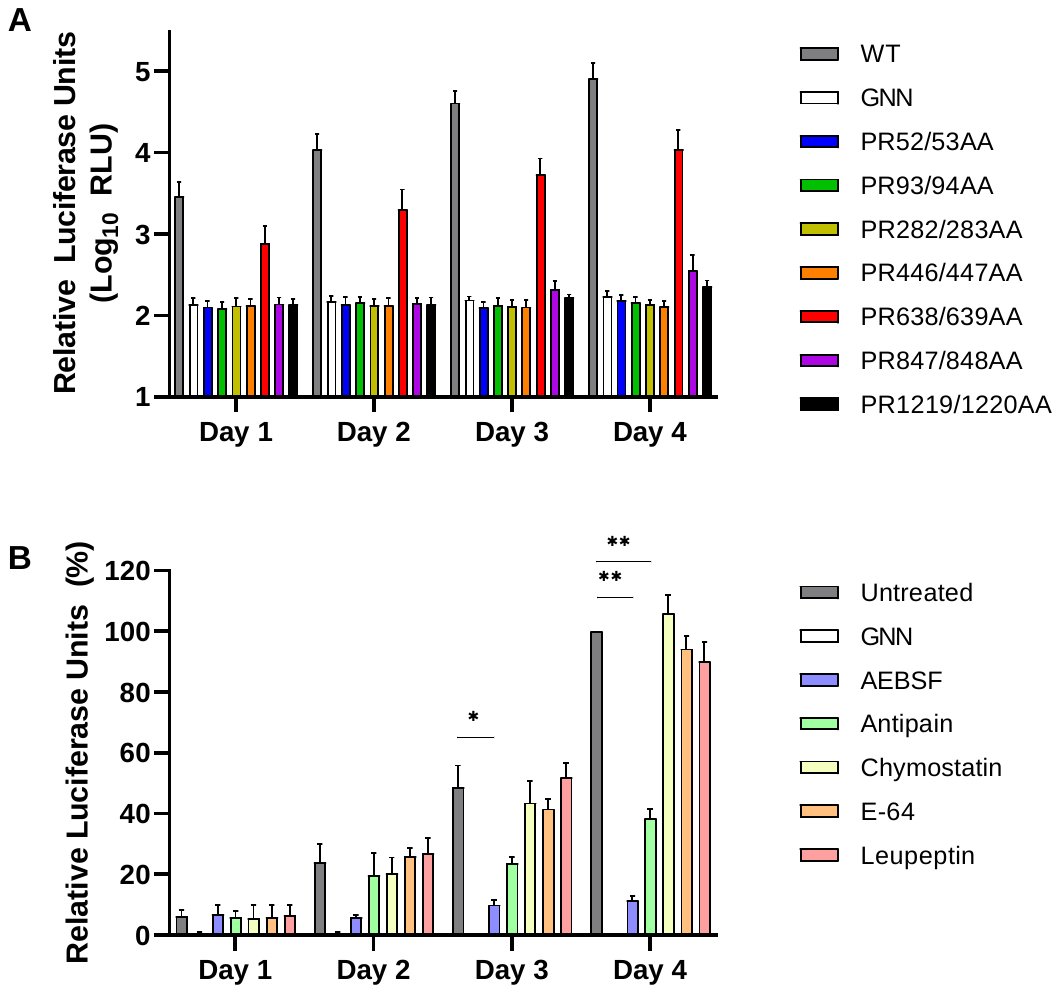

**Figure S2. Preventing thrombin proteolysis prevents HEV replication. (A)** Huh7 cells were electroporated with HEV replicon RNA containing the indicated mutations at predicted thrombin cleavage junctions, in addition to the WT and GNN control replicons. Cells were harvested at the indicated times post-electroporation and luciferase activity determined. Data shown represents log_10_ of mean relative luciferase activity (n = 3 +/- SEM). **(B)** Huh7 cells were electroporated with the WT HEV replicon RNA or GNN control replicon before the addition of AEBSF (2.5 µg / mL), antipain (50 µg / mL), chymostatin (25 µg / mL), E-64 (100 µg / mL) or leupeptin (175 µg / mL) 24 h post electroporation. Cells were harvested at the indicated times post-electroporation and luciferase activity determined. Data shown represents mean relative luciferase activity (n = 3 +/- SEM, *=p<0.05, **=p<0.01 compared to WT).
